# Using citizen science monitoring data in species distribution models to inform isotopic assignment of migratory connectivity in wetland birds

**DOI:** 10.1101/144527

**Authors:** Auriel M. V. Fournier, Kiel L. Drake, Douglas C. Tozer

## Abstract

Stable isotopes have been used to estimate migratory connectivity in many species. Estimates are often greatly improved when coupled with species distribution models (SDMs), which temper estimates in relation to occurrence. SDMs can be constructed using from point locality data from a variety of sources including extensive monitoring data typically collected by citizen scientists. However, one potential issue with SDM is that these data oven have sampling bias. To avoid this potential bias, an approach using SDMs based on marsh bird monitoring program data collected by citizen scientists and other participants following protocols specifically designed to maximize detections of species of interest at locations representative of the species range. We then used the SDMs to refine isotopic assignments of breeding areas of autumn-migrating and wintering Sora *(Porzana Carolina)*, Virginia Rails *(Rallus limicola)*, and Yellow Rails *(Coturnicops noveboracensis)* based on feathers collected from individuals caught at various locations in the United States from Minnesota south to Louisiana and South Carolina. Sora were assigned to an area that included much of the western U.S. and prairie Canada, covering parts of the Pacific, Central, and Mississippi Flyways. Yellow Rails were assigned to a broad area along Hudson and James Bay in northern Manitoba and Ontario, as well as smaller parts of Quebec, Minnesota, Wisconsin, and Michigan, including parts of the Mississippi and Atlantic Flyways. Virginia Rails were from several discrete areas, including parts of Colorado, New Mexico, the central valley of California, and southern Saskatchewan and Manitoba in the Pacific and Central Flyways. Our study demonstrates extensive data from organized citizen science monitoring programs are especially useful for improving isotopic assignments of migratory connectivity in birds, which can ultimately lead to better informed management decisions and conservation actions.

## Introduction

Determining links among breeding, migratory stopover, and wintering areas for different populations of migratory birds (hereafter ‘migratory connectivity’) is critically important for conserving species throughout their annual cycle (Webster et al. 2002, Hobson et al. 2014). Knowledge on migratory connectivity informs full life cycle conservation by associating populations with areas that they inhabit. Connecting populations and habitat is essential to identifying limiting factors and can permit more effective responses to threats by targeting the most affect parts of populations or the annual cycle (Norris and Taylor 2006, Taylor and Norris 2010, Rushing et al. 2016). Most studies examine migratory connectivity between wintering and breeding areas, but connectivity with stopover habitat during migration is also important in understanding factors that limit populations throughout their annual cycle (e.g., Hobson et al. 2014; 2015).

There are a variety of effective methods for estimating migratory connectivity of birds including mark-recapture (Ryder et al. 2011), archival biologgers (Ryder et al. 2011, Salewski et al. 2013, Hallworth et al. 2013), collaborative radio tracking networks (Taylor et al. 2017), and satellite transmitters (Krementz et al. 2011). Despite a diversity of methods, these techniques will not work with all species. When a transmitter is too heavy relative to the weight of the species it is unsafe to attach the device. When a species has low site fidelity among years it becomes impractical to relocate and recapture individuals to retrieve the archival biologgers and the data the devices contain. In addition, collaborative radio-tracking networks, although extremely promising in the near-future for broad scale studies, are currently unavailable in most areas. In situations when the above approaches are ineffective, isotopes can be used because individuals need to be captured only once to obtain samples (e.g., feathers, toenails, blood), and no tracking devices need to be attached. In North America, the ratio of hydrogen isotopes (δ^2^H) follows a spatial gradient from northwest to southeast and has been widely used to examine migratory connectivity of many species (Hobson and Wassenar 2008, Guillemain et al. 2014a, Butler et al. 2016). One disadvantage of stable hydrogen isotopes is the resulting coarse geographic assignments, which can limit the level of inference, but even this limited inference can inform conservation if it’s the only information available (Hobson and Wassenaar 2008). The incorporation of additional data, such as environmental variables, genetic information, band recovery data, and predictions from species distribution models (SDMs), can improve geographic assignment because populations are not equally spread over space (Royle and Rubenstein 2004, Hobson et al. 2013, Rushing et al. 2014, Ruegg et al. 2016). The results lead to assignments that are more informative for conservation and management purposes (Haig et al. 1998, Webster et al. 2002, Hobson 2005).

Of the many options for refining isotopic assignments of migratory connectivty, SDMs show excellent utility. The models can be used to predict species occurrence or abundance across vast unsampled areas, often with reasonable precision and accuracy based on existing data (Elith and Leathwick 2009). This information can then be coupled with isotopic assignments to produce refinements in relation to species occurrence or abundance. The most useful SDMs for this purpose are ones based on extensive representative datasets through space and time.

Species abundance data at broad scales are easily obtained from citizen science monitoring programs. These programs operate by engaging volunteers and training them to follow standardized survey protocols to collect reliable monitoring data. Due to the volunteer nature of the programs combined with widespread engagement of participants, they often produce large sample sizes over broad spatial distributions that benefit SDM development. For instance, Fournier et al. (2016) used haphazard presence-only citizen science data from eBird (Sullivan et al. 2009) to refine stable isotope assignments of migratory connectivity in the Virginia Rail *(Rallus limicola)*. The approach was successful and substaintially improved the refinement of isotopic assignments. However, the authors noted challenges due to potential biases caused by factors driving where and how observers conduct surveys.

One way to overcome perceived bias associated with haphazard presence-only data is to use data collected by formal monitoring programs. In these programs, participants collect data at locations regardless of whether the species of interest was detected or not, following established protocols designed to maximize detection probability (e.g. Conway 2011). Organized monitoring programs typically collect data at pre-determined randomly-chosen survey locations, making the data representative of entire populations. Therefore, data from organized bird monitoring programs is more suitable than presence only data for developing SDMs to refine istopic assignments of migratory conenctivity.

In this paper, we demonstrate the use of SDMs to refine isotopic assignments of migratory connectivity in Sora *(Porzana carolina)*, Virginia Rail, and Yellow Rail *(Coturnicops noveboracensis)* using data collected by citizen scientists and other participants in organized marsh bird monitoring programs and feathers collected from individuals caught at autumn migration and wintering locations in the United States ranging from Minnesota to Louisiana and South Carolina. We chose these three rail species, in part, because they are elusive wetland birds that breed across a wide swath of North America, but are poorly studied (Eddleman et al. 1988). The species are of concern because they stopover in highly modified landscapes where wetland loss ranges 60-90%, and their populations are thought to be declining, but are not clearly understood (Reid 1989, Case and McCool 2009, Ducks Unlimited Canada 2010, Dahl 2011). In addition, the Sora and Virginia Rail are game bird species in some jurisdictions (Tacha and Braun 1994), while the Yellow Rail is a species of special concern in Canada (Alvo and Robert 2009). Knowledge of migratory connectivity in these three rail species is only now beginning to emerge (Butler et al. 2016, Fournier et al. 2016), and is needed to inform conservation and management efforts. Studying broad scale migratory connectivity in the three species is also currently unsuitable with any of the methods listed above, except isotopes. Together, these characteristics made the species worthy candidates with which to demonstrate the method and approach.

## Methods

### Field

#### Migrating and Wintering Individuals

Sora, Yellow Rail, and Virginia Rail were captured using dipnets from all-terrain vehicles during autumn migration between August and October 2015 at 10 sites in Missouri, USA (Perkins et al. 2010). Sora, Virginia Rail and Yellow Rail feathers from other migratory locations (Minnesota, Michigan, South Carolina, Ohio, and Arkansas, USA) and wintering locations (Louisiana and Mississippi, USA) were collected opportunistically by hunters and researchers from August through December 2015 (Table S2). The first primary feather, which is grown on the breeding grounds (Pyle 2008) and therefore has the isotopic signature of that location, was removed from each individual. Previously collected Yellow Rail feathers from another project were included to increase sample size and these feathers were collected in Missouri during autumn migration in 2013 and 2014.

#### Breeding Individuals

Sora, Yellow Rail and Virginia Rail were captured on foot during night using call broadcast lures and a dipnet in late-June and July 2015 near Foam Lake, Saskatchewan, Canada (51.6601, -103.5538). Captures began at dusk and ran until dawn. Similar to migrants, the first primary feather was removed from each individual.

### Laboratory

Feathers were cleaned with phosphate-free detergent and 2:1 chloroform methanol solution, rinsed them in deionized water, and dried them at 50 °C for 24 hours. A total of 0.350 mg of material was weighed into silver capsules (Elemental Microanalysis, part# d2302) and analyzed by coupled pyrolysis/isotope-ratio mass spectrometry using a thermo-chemical elemental analyzer (TC/EA) (Thermo Scientific) interfaced to a Thermo Scientific Delta V Plus configured through a CONFLO IV for automated continuous flow gas-isotope ratio mass spectrometer (CF-IRMS) at the Colorado Plateau Stable Isotope Laboratory at Northern Arizona University.

Given that ~20% of the δ^2^H in feathers exchanges freely with ambient water vapor (Wassenaar & Hobson 2003), we analyzed feathers concurrently with three calibrated keratin standards (Keratin – SC Lot SJ (powdered) mean = –120.7 ± 1.1 ‰, expected = –121.6 ‰, n=32; CBS – caribou hoof (powdered) mean= –198.5 ± 1.1 ‰, expected = –197.0 ‰, n=10; KHS – Kudo horn (powdered) mean= –55.1 ± 1.0 ‰, expected = –54.1 ‰, n=10) to allow for future comparison across laboratories (Wassenaar & Hobson 2003). We report the non-exchangeable δ^2^H fraction in parts per mil (‰) normalized to the Vienna Standard Mean Ocean Water-Standard Light Antarctic Precipitation (VSMOW-SLAP) standard.

### Species Distribution Models

We used count data from 7,146 100-m-radius plots surveyed largely by citizen scientists and other participants in Bird Studies Canada’s Great Lakes, Québec, and Prairie marsh monitoring programs (Bird Studies Canada 2017a; Tozer 2013, 2016) available through Nature Counts (Bird Studies Canada 2017b), and by observers in the North American Marsh Bird Monitoring Program at various National Wildlife Refuges available from the Midwest Avian Data Center (Figure 1; Koch et al. 2010) to construct SDMs. These data spanned 1995-2015 and were collected under a slightly modified version of the Standardized North American Marsh Bird Monitoring Protocol (e.g., Tozer et al. 2016), which included the use of standardized call broadcasts of Sora, Yellow Rail, and Virginia Rail during point counts to increase detection probability (Conway 2011). We collapsed the dataset so the response was the highest count at each point across all years. We did this instead of formally modeling detection probability to keep the models simple and more manageable, and to overcome challenges with modeling detection across points surveyed in different years and in different numbers of years. Selecting the maximum count across all years surveyed gives a conservative estimate of abundance of each species at each point. This yielded 929 Sora, 695 Virginia Rail, 39 Yellow Rail points where at least one individual was detected and 4,056 Sora, 4,290 Virginia Rail, and 4,946 Yellow Rail points were each species was not detected in any year.

**Figure 1.**
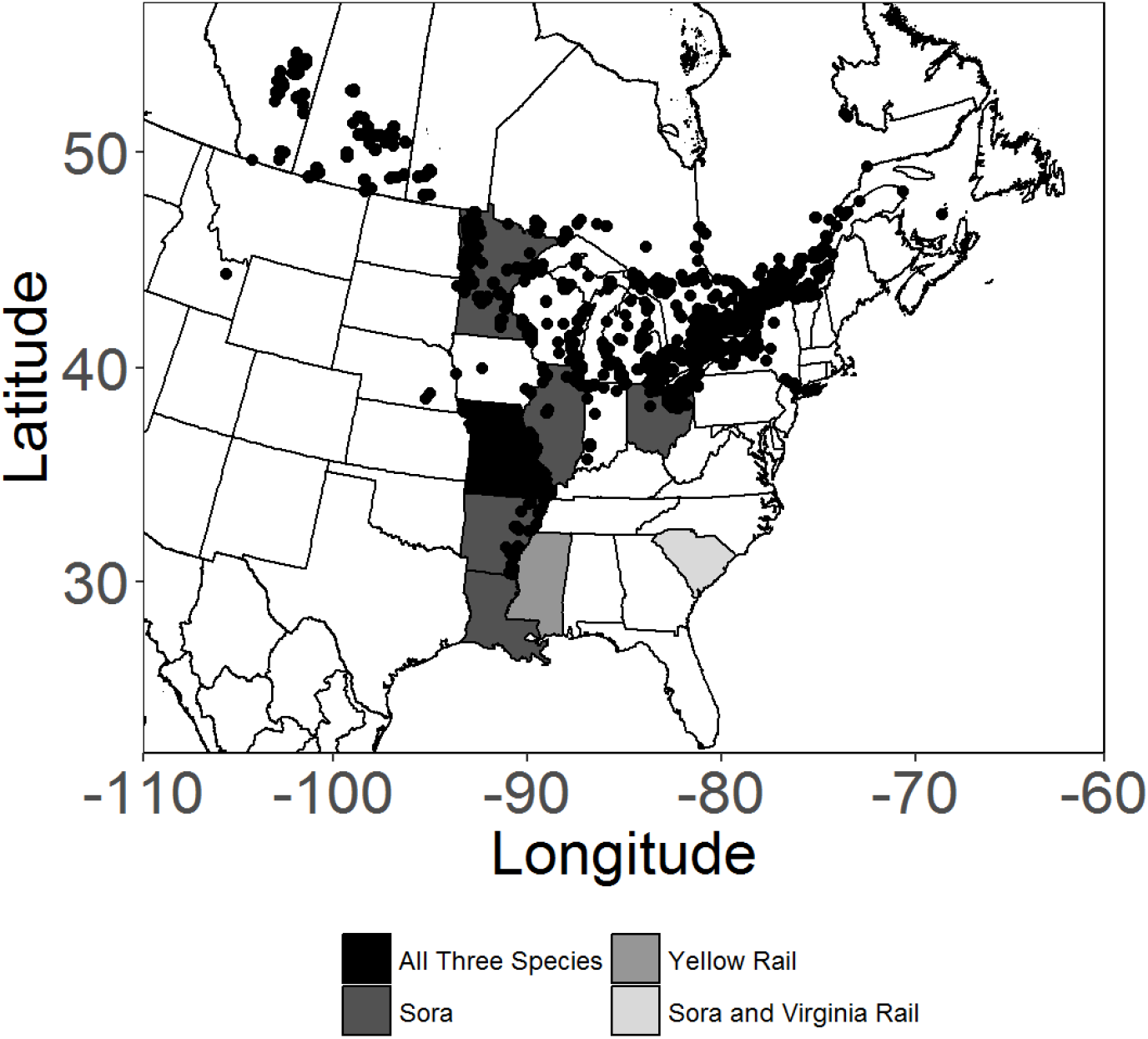
Locations of marsh bird monitoring program survey points used to develop species distribution models, and states where autumn-migrating and wintering rails were captured and sampled or isotopic analysis.

We created species distribution models describing Sora, Yellow Rail and Virginia Rail abundance using 11 raster layers (1 km^2^ resolution) representing land cover (Latifovic et al. 2002), wetland presence (Lehner & Döll 2004), and bioclimatic parameters. We chose these layers because they likely influence precipitation, and thus stable isotope ratios across North America, as well as the distribution and abundance of the three species we considered. We removed variables which were correlated (Pearson’s correlation coefficient >=75%, See Table S1). We constrained predictions to each species’ summer range (BirdLife International & NatureServe 2015). Within each species’ summer range, our goal, similar to Fournier et al. (2016), was to generate a SDM with the greatest predictive accuracy (see details below), but not necessarily informative for inferring environmental relationships (Merow, Smith and Silander 2013). We included all 11 environmental covariates in the analysis and used leave-one-out jackknifing to identify covariates that reduced the predictive power of the model, which were then removed. All modeling took place in R (R Core Team 2016, version 3.3.2).

#### Spatially explicit assignment of geographic origins

We used the methods and code of Van Wilgenburg and Hobson 2011 to perform our spatially explicit isotopic assignments for each individual. Below is a summary of those methods. We used a likelihood-based assignment that incorporated estimates of uncertainty (Royle and Rubenstein 2004). Expected δ^2^Hfeather values were calculated by regressing raw δ^2^H feather values of sampled feathers on mean annual growing season δ^2^H in precipitation at the site of collection. This calibration was necessary to account for systematic differences between the δ^2^H of sampled feathers and δ^2^H in precipitation. Because we only had feathers from one breeding ground location, we included data from other projects in our linear regression of δ^2^H of flight feathers to mean annual growing season δ^2^H across North America (~37 × 37 km resolution; Bowen et al. 2005). This known-origin dataset included feathers from Foam Lake Saskatchewan (45 Sora feathers, 30 Yellow Rail Feathers and 4 Virginia Rail), and 10 Virginia Rail feathers from one location from Fournier et al. (2016), along with 44 King Rail feathers from Perkins (2007), including 13 museum specimens from 11 different localities and 31 live captured King Rail specimens. In total we had 133 feathers from 14 different localities (for additional detail on the feathers from locations outside of Saskatchewan see Appendix S1 in Fournier et al. (2016)). Because of small sample size for Yellow Rails in 2015, we also included feathers from autumn migration in 2013 and 2014. We did not find a significant difference between the median δ^2^H values in Yellow Rails among years (ANOVA F = 0.11, df = 21, p = 0.91; Figure 2), suggesting that inter-annual variability in feather δ^2^H was unlikely to be a significant source of variation for our analysis so we combined annual samples. We regressed our data of known-origin feathers against δ^2^H precipitation to derive the calibration equation (δ^2^H_corrected_ = -52.36 + 0.83[δ^2^H_precipitation_]).

**Figure 2.**
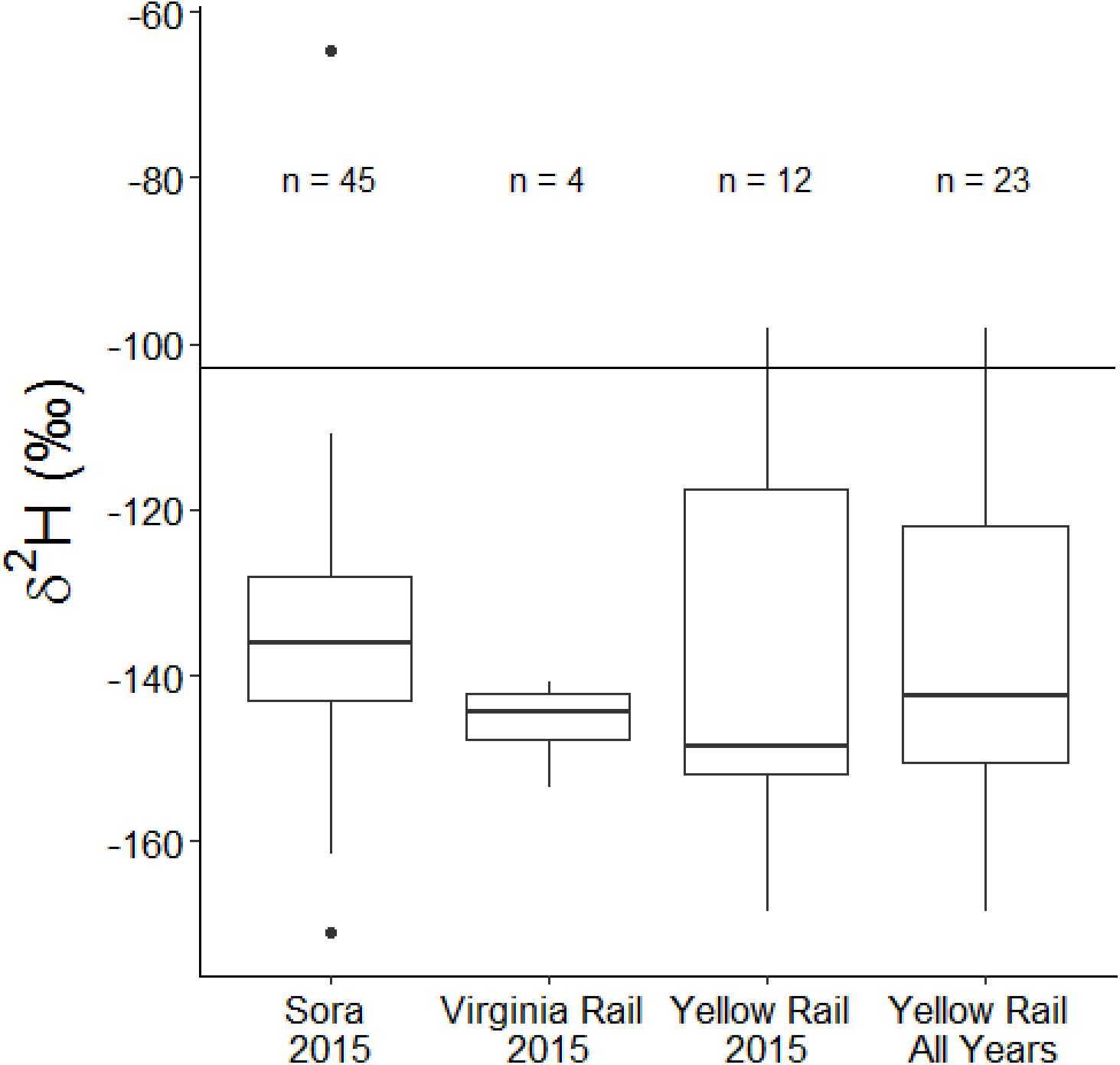
– Distribution of *δ*^2^H values of feathers from rails caught at Foam Lake, Saskatchewan, Canada. The horizontal line represents *δ*^2^H in precipitation from Bowen et al. (2005).

For each feather we assessed the probability that any cell within the expected values was the origin of that individual using a normal probability density function as follows:

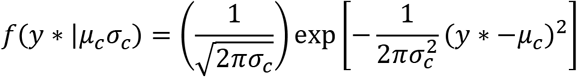

Where *f*(*y* * |*μ_c_σ_c_*) represents the probability that a given cell (c) within the δ^2^HF isoscape represents a potential origin for an individual of unknown origin (y*), given the expected mean δ^2^HF for that cell (*μ_c_*) from the calibrated δ^2^HF isoscape and the expected standard deviation (*σ_c_*) of δ^2^HF between individuals growing their feathers at the same locality. To assign probable breeding areas to samples within a particular state, we summed the assignments from each feather sample in units of the number of rails with origins consistent with a given pixel and converted to proportions to enable comparisons with other states, which we report only in the supplementary material. For each individual we produced a surface of spatially explicit probability densities (i.e., one surface per bird in a sample). We then incorporated the prior probabilities from our SDM by applying Bayes’s Rule (Van Wilgenburg and Hobson 2011). To depict these origins across the entire sample size we assigned each feather to the base map individually by determining the odds that any given assigned origin was correct relative to the odds it was incorrect. Based on 3:1 odds that a given bird had originated from within the range we recorded the set of raster cells that defined the upper 75% of estimated origins and coded them as 1, all others as 0. We choose 3:1 odds based on Van Wilgenburg and Hobson (2011) where this ratio provided a compromise between the possibility of being incorrect and the bird assignment geographic resolution. The results of the individual assignments were then summed over all individuals, by addition of the surfaces. We facilitated this step by rescaling the posterior probabilities (f_x_) relative to the maximum value within the posterior probability surface prior to applying the odds-ratio-based reclassification.

We made assignments using functions within the R statistical computing environment (R Core Team 2016, version 3.3.2) using the ‘raster’ package (Hijmans 2016, version 2.5-8). To make our results even more relevant for conservation, we also visually inspected assignments to determine broad overlap with traditional flyways used as administrative units by migratory game bird managers (US Fish and Wildlife Service 2017).

## Results

### Captures

We captured 142 southbound autumn migrating and wintering rails across the southern U.S., and 79 breeding rails at a wetland complex in Saskatchewan, Canada. Sora comprised the bulk of migrant and wintering samples (88%; 117 individuals; 8 states) followed by Virginia Rails (7%; 9 individuals; 2 states) and Yellow Rails (5%; 11 individuals; 2 states). Sora also comprised the bulk of breeder samples (57%; 45 individuals) followed by Yellow Rails (38%; 30 individuals) and Virginia Rails (5%, 4 individuals). See Table S2 for more details.

### Species Distribution Models

All three species distribution models fit the data (Homer-Lemeshow Goodness of Fit Test, Sora *χ*2 = 4.7, df = 8, p = 0.7; Virginia Rail *χ*2= 4.7, df = 8, p = 0.7; Yellow Rail *χ*2= 4.4, df = 8 p = 0.8). The top species distribution model for Yellow Rail contained mean temperature of driest quarter (β = -0.30, SE = 0.09, p = 0.002), mean temperature of warmest quarter (β = 0.28, SE = 0.14, p = 0.008), mean diurnal range (mean of monthly (max temp-min temp)) (β = 0.60, SE = 0.16, p <0.001) and a significant interaction between latitude and longitude (β = 2.09, SE = 0.59, p <0.001). The top species distribution model for Virginia Rail contained temperature seasonality (β = -0.002, SE = 0.0007, p = 0.002). The top species distribution model for Sora included annual mean temperature (β = 0.07, SE = 0.02, p=0.003), mean temperature of the warmest quarter (β = -0.09, SE = 0.02, p<0.001) and temperature seasonality (β = 0.001, SE = 0.0002, p <0.001).

### Isotopic Assignments

Sora were assigned to an area that included much of the western U.S. and prairie Canada, covering parts of the Pacific, Central, and Mississippi flyways (Figure 3). Yellow Rails were assigned to a broad area along Hudson and James Bay in northern Manitoba and Ontario, as well as smaller parts of Quebec, Minnesota, Wisconsin, and Michigan, including parts of the Mississippi and Atlantic flyways (Figure 3). Virginia Rails were from several discrete areas, including the southern part of their breeding range in parts of Colorado, New Mexico, the central valley of California, and southern Saskatchewan and Manitoba in the Pacific and Central flyways (Figure 3). Due to small sample size (Table S2), we do not include comparison of breeding ground assignments among states, although for the interested reader we include maps of these assignments in the supplementary material (Figure S2, S3, S4).

**Figure 3.**
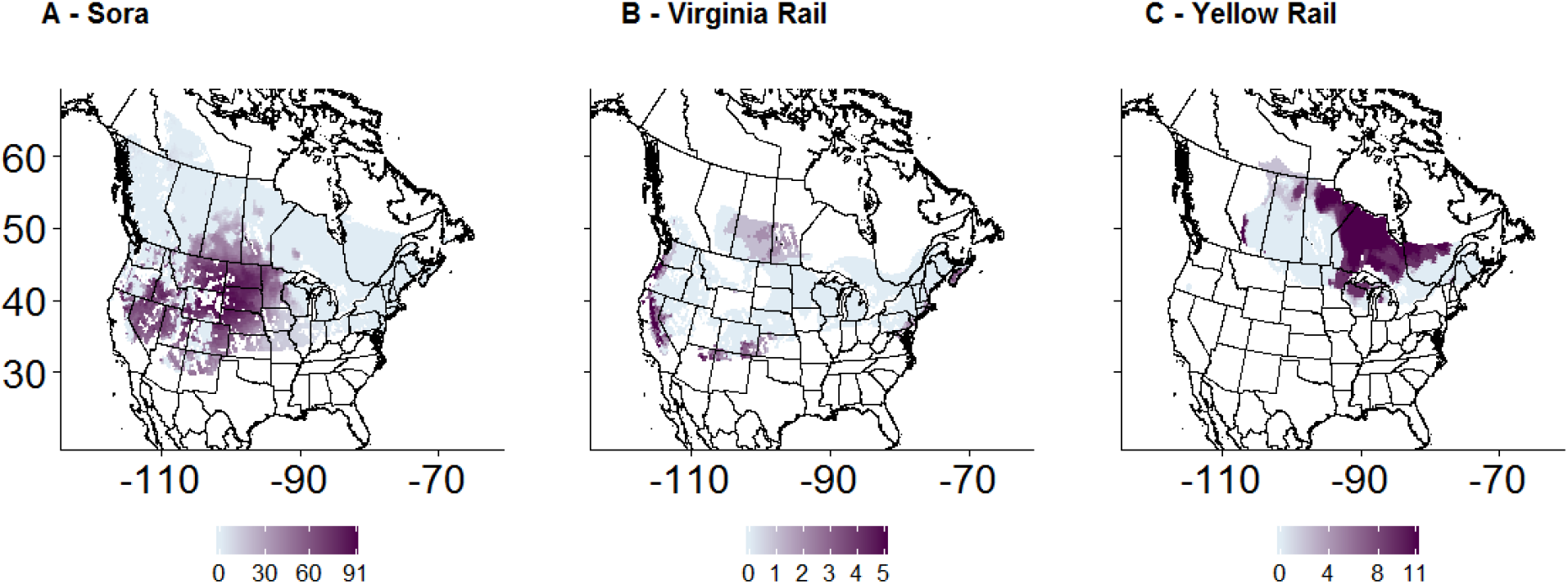
– Cumulative assignment of breeding areas of autumn migrating and wintering Sora *(Porzana carolina)*, Virginia Rails *(Rallus limicola)* and Yellow Rails *(Corturnicops noveboracensis)* based on expected *δ*^2^H_feather_ values using regional monitoring data in a species distribution model as an informative prior. Each individual bird’s assignment surface represents the area where the bird is like from with 3:1 odds and then those surfaces are summed to form the cumulative assignment for all individuals from that species.

## Discussion

We demonstrated the use of SDMs to inform isotopic assignments of migratory connectivity in wetland birds, based on organized marsh bird monitoring program data collected by citizen scientists and other participants. We found these data to be especially useful for this purpose for reasons related to sample size, search effort, detection probability, and ease of obtaining data, which we elaborate further below.

Use of SDMs to refine isotopic assignments of migratory connectivity should be based on extensive datasets through space and time. Such data are most easily obtained by researchers from citizen science monitoring programs. These programs normally involve careful training of participants to follow well-established and tested field protocols that produce reliable data. They also typically engage impressive numbers of participants to survey numerous locations throughout large portions of the range of occurrence of species of interest. These characteristics produce datasets with large sample sizes that are amenable for capturing the range of conditions and circumstances under which species occur, leading to better predictions based on SDMs for refining isotopic assignments of migratory connectivity than from isotopes alone.

Various extensive citizen science datasets suitable for SDM deveopment are freely-available to researchers. Most of these and other useful sources of data are easily obtained through the various information nodes of the Avian Knowledge Network (2017), such as the ones used to obtain data for this paper: Nature Counts (Bird Studies Canada 2017b) and Midwest Avian Data Center (Koch et al. 2010).

Some available datasets, however, are more useful or easier to implement than others for developing SDMs. Like the programs that produced data for this paper, some monitoring programs pre-select survey locations so they are representative of larger areas of inference, typically by using various randomization procedures (e.g., Johnson et al. 2009). Many of these programs also record data regardless of whether certain species were detected or not, following protocols specifically designed to maximize detections of species of interest (e.g., Conway 2011). Such protocols include restrictions on the time of day and season, type of weather, and the amount of background noise that is acceptable during surveys (e.g., Tozer et al. 2016). They also include requirements on the minimum number of visits per survey location, and the total duration of each survey, plus some use standardized call broadcasts to increase the probability of detection of especially elusive species. All of these characteristics provide more reliable information on the presence or absence or abundance of species of interest at a particular point. These programs, which are dedicated to generating reliable, representative data on occurrences or counts of species across specific areas of inference may be the best choice, when available, for developing SDMs to refine isotopic assignments of migratory connectivity.

By contrast, monitoring programs that lack the standardized restrictions and guidelines noted above can pose challenges for SDM development. This was shown by the additional bias-correction analysis that Fournier et al. (2016) were required to perform during their use of SDMs based on eBird data to refine isotopic assignments of migratory connectivity in the Virginia Rail. The bias was thought to occur because there are no restrictions on where and how eBird participants observe rails or other species, and because of the nature of eBird data it did not meet the assumption that presence data points were randomly distributed (Yackulick et al. 2012). While this flexibility is a major advantage of eBird and other programs like it for numerous other applications, the lack of organized standardization of surveys can cause challenges for SDM development (Yackulic et al 2012).

Our SDMs might have provided better assignment resolution if finer-scale habitat covariates, especially wetland cover, were available in a consistent format across Canada and the U.S. Our analysis might also have been improved by simultaneously considering another isotope, such as Sulphur (δ^34^S). Some rails use brackish or saline habitats during the breeding season, and this would be reflected in their δ^34^S feather signatures, potentially helping to further refine assignments (Hobson et al. 2012, Butler et al. 2016). The incorporation of genetic information could have been beneficial, though currently, to our knowledge, such information is not available for rails.

We combined isotopic signatures of the largest sample of autumn-migrating and wintering rails with SDMs based on organized marsh bird monitoring data to produce the most extensive estimates of migratory connectivity of three rail species currently available. We found that the migratory connectivity of the three species included wide-ranging breeding areas, including more than one migratory game bird flyway in the two hunted species—results useful for improving conservation of these poorly-studied species—although additional work is needed to fully establish patterns. Extensive data from organized citizen science monitoring programs are especially useful for improving isotopic assignments of migratory connectivity in birds, which will ultimately lead to better management and conservation of species.

## Acknowledgements

Funding provided through the Arkansas Cooperative Fish and Wildlife Research Unit, Bird Studies Canada’s Long Point Waterfowl and Wetlands Research Program, Garden Club of America’s Frances M. Peacock Scholarship for Native Bird Habitat, SC Johnson, and The Bluff’s Hunting Club. A special thank you goes to the many volunteers and employees who have contributed to provincial and state marsh bird monitoring programs. Thanks to Matt Boone, Nick Seeger, Dan Datlof, Dan Holm, Hailee Pavisich, Patrick Turgeon, David Anderson, Erik Johnson, Christine Hand, John Simpson and LeeAnn Latremouille, Avian Events Support Team’s Yellow Rails and Rice Festival, Louisiana Bird Observatory, and Audubon Louisiana who helped capture birds and collect feathers for this project. Special thanks go to Alex Bond and David Krementz for comments that greatly improved an earlier version of the manuscript. Feathers were collected in Canada under federal bird banding permit #10842, #10842C, #10842D, and #10842F, and in the U.S. under federal bird banding permit #23002. University of Arkansas IACUC protocols #15049, #15023, and #13044 covered this project.

## References

Alvo, R. and Robert, M. 1999. COSEWIC status report on the Yellow Rail *Coturnicops noveboracensis* in Canada. Committee on the Status of Endangered Wildlife in Canada. Ottawa. 1–62 pp. http://www.registrelep-sararegistry.gc.ca/virtual_sara/files/cosewic/sr_Yellow%20Rail_0810_e.pdf (Accessed 13 September 2016)

Bird Studies Canada. 2017a. Marsh Monitoring Program. http://www.birdscanada.org/volunteer/natmmp/index.jsp?lang=EN (Accessed 25 January 2017).

Bird Studies Canada. 2017b. Nature Counts. http://www.birdscanada.org/birdmon/default/main.jsp (Accessed 5 January 2017).

Bird Studies Canada and Cornel Lab of Ornithology. 2017. Project FeederWatch. http://feederwatch.org/ (Accessed 25 January 2017).

BirdLife International and NatureServe. 2015. Bird species distribution maps of the world. BirdLife International, Cambridge, UK and NatureServe, Arlington, USA.

Butler, C. J., Wilson, J.K., Frazee, S. R., and Kelly, J. F. 2016. A comparison of the origins of yellow rails *(Coturnicops noveboracensis)* wintering in Oklahoma and Texas, USA. – Waterbirds. 39: 156–164.

Bowen, G.J. 2008. Spatial analysis of the intra-annual variation of precipitation isotope ratios and its climatological corollaries - J. Geophys. Res. Atmos. 113: D05113. doi:10.1029/2007JD009295

Bowen, G. J., Wassenaar, L.I. and Hobson, K.A. 2005. Global application of stable hydrogen and oxygen isotopes to wildlife forensics. - Oecologia. 143: 337–348

Case, D. J., and McCool, D. D. 2009. Priority information needs for rails and snipe.

Conway, C. J. 2011. Standardized North American marsh bird monitoring protocol. – Waterbirds. 34: 319–346.

Dahl, T. E. 2011. Status and trends of wetlands in the conterminous United States 2004 to 2009. US Department of the Interior, US Fish and Wildlife Service, Fisheries and Habitat Conservation.

Ducks Unlimited Canada. 2010. Southern Ontario wetland conversion analysis. Final report, March 2010. Published by Ducks Unlimited Canada, Barrie, ON. http://www.ducks.ca/assets/2010/10/duc_ontariowca_optimized.pdf (Accessed 5 June 2015).

Eddleman, W. R., F. L. Knopf, B. Meanley, F. A. Reid, and R. Zembal. 1988. Conservation of North American Rallids. - The Wilson Bulletin 100:458–475.

Elith, J., and Leathwick, J. R. 2009. Species Distribution Models: Ecological Explanation and Prediction across Space and Time.

Fournier, A. M. V., Sullivan, A. R., Bump, J. K., Perkins, M., Shieldcastle, M. C. and King, S. L. 2016. Combining citizen science species distribution models and stable isotopes reveals migratory connectivity in the secretive Virginia rail. J Appl Ecol. doi: 10.1111/1365-2664.12723

Guillemain, M., Van Wilgenburg, S. L., Legagneaux, P. and Hobson, K. A. 2014a. Assessing geographic origins of Teal (Anas crecca) through stable-hydrogen isotope analyses of feathers and ring-recoveries. - J. Ornithol. 155: 165–172.

Haig, S. M., Mehlman, D. W. and Oring, L. W. 1998. Avian movements and wetland connectivity in landscape conservation. - Conserv. Biol. 12: 749–758.

Hallworth, M. T., Studds, C. E., Sillett, T. S. and Marra, P. P. 2013. Do archival light-level geolocators and stable hydrogen isotopes provide comparable estimates of breeding-ground origin? – Auk. 130: 273–282.

Hijmans, R.J. 2016. raster: Geographic Data Analysis and Modeling. R package version 2.5–8. https://CRAN.R-project.org/package=raster

Hobson, K. A. 2005. Using stable isotopes to trace long distance dispersal in birds and other taxa. - Divers. Distrib. 11: 157–164.

Hobson, K. A., and Wassenaar, L. I. 2008. Tracking animal migration with stable isotopes. Elsevier.

Hobson, K. A., Van Wilgenburg, S. L., Wassenaar, L. I., Powell, R. L., Still, C. J., and Craine, J. M. 2012. A multi-isotope (δ^2^H, δ^13^C, δ^15^N) approach to establishing migratory connectivity in Palearctic Afrotropical migrants: An example using Wood Warblers Phylloscopus sibilatrix. - Ecosphere, 3: 44.

Hobson, K. A., Van Wilgenburg, S. L., Ferrand, Y., Gossman, F. and Bastat, C. 2013. A stable isotope (^2^H) approach to deriving origins of harvested woodcock taken in France. - Eur. J. Wildl. Res. 59: 881–892.

Hobson, K. A., Van Wilgenburg, S. L., Faaborg, J., Toms J.D., Rengifo, C., Sosa, A. L., Aubry, Y., Aguilar, R. B. 2014. Connecting breeding and wintering grounds of Neotropical migrant songbirds using stable hydrogen isotopes: a call for an isotopic atlas of migratory connectivity. - J Field Ornithol. 85:237–257.

Hobson, K. A., Van Wilgenburg, S. L., Dunn, E. H., Hussell, D. J. T., Taylor, P. D., and Collister, D. M. 2015. Predicting origins of passerines migrating through Canadian migration monitoring stations using stable hydrogen isotope analyses of feathers: a new tool for bird conservation. - Avian Conserve Ecol 10: 3. http://dx.doi.org/10.5751/ACE-00719-100103.

Koch, K., D. Moody, S. Michaile, M. Magana, M. Fitzgibbon, G. Rowell, T. Will, and G. Ballard. 2010. The Midwest Avian Data Center. [web application]. Petaluma, California. http://data.pointblue.org/partners/mwadc.

Krementz, D. G., Asante, K. and Naylor, L. W. 2011. Spring migration of mallards from Arkansas as determined by satellite telemetry. - J. Fish Wildl. Manag. 2: 156–168.

Latifovic, R., Zhu, Z.-L., Cihlar, J. & Giri, C. 2002. Land cover of North America 2000. Natural Resources Canada, Canada Center for Remote Sensing, US Geological Service EROS Data Center

Lehner, B. and Döll, P. 2004. Development and validation of a global database of lakes, reservoirs and wetlands. - J. Hydro. 296: 1–22.

Link, W.A. & Sauer, J.R. 1998. Estimating population change from count data: Application to the North American Breeding Bird Survey. Ecological Applications, 8, 258–268.

Merow, C., Smith, M. J. and Silander, J. A. 2013. A practical guide to MaxEnt for modeling species’ distributions: what it does, and why inputs and settings matter. Ecography, 36: 1058–1069. doi: 10.1111/j.1600-0587.2013.07872.x

National Audubon Society 2010. The Christmas Bird Count Historical Results [Online]. Available http://www.christmasbirdcount.org

Norris, D. R. and Taylor, C. M. 2006. Predicting the consequences of carry-over effects for migratory populations. – Biol. Letters 22:148–151

Perkins, M. 2007. The Use of Stable Isotopes To Determine The Ratio Of Resident to Migrant King Rails In Southern Louisana and Texas. Louisana State University.

Perkins, M., King, S. L. and Linscombe, J. 2010. Effectiveness of capture techniques for rails in emergent marsh and agricultural wetlands. - Waterbirds 33: 376–380.

Price, J.T., Droege, S. & Price, A. 1995. Summer atlas of North American birds. Academic Press, London.

Pyle, P. 2008. Identification Guide to North American Birds Part II (First Edit). Point Reyes Station, California: Slate Creek Press.

R Core Team. 2016. R: A language and environment for statistical computing. R Foundation for Statistical Computing, Vienna, Austria. Version 3.3.2

Reid, F. A. 1989. Differential habitat use by waterbirds in a managed wetland complex. University of Missouri-Columbia.

Royle, J. A., and Rubenstein, D. R. 2004. The role of species abundance in determining breeding origins of migratory birds with stable isotopes. – Eco. Apps. 14: 1780–1788.

Ruegg K.C., Anderson E., Harrigan R. J., Paxton K. L., Kelly J., Moore F., Smith T. B. 2016. bioRxiv 085456; doi: 10.1101/085456

Rushing, C. S., Ryder, T. B., Saracco, J. F. and Marra, P. P. 2014. Assessing migratory connectivity for a long-distance migratory bird using multiple intrinsic markers. Ecological Applications, 24: 445–456. doi: 10.1890/13-1091.1

Rushing, C. S., Ryder, T. B., Mara, P. P. 2016. Quantifying drivers of population dynamics for a migratory bird throughout the annual cycle. - Proc. R. Soc. B 283:20152846.

Ryder, T. B., J. W. Fox, and P. P. Marra. 2011. Estimating Migratory Connectivity of Gray Catbirds (Dumetella carolinensis) using Geolocator and Mark—Recapture Data. The Auk 128:448–453.

Salewski, V., Flade, M., Poluda, A., Kiljan, G., Liechti, F., Lisovski, S. and Hahn, S. 2013. An unknown migration route of the “globally threatened” Aquatic Warbler revealed by geolocators. - J. Ornithol. 154: 549–552.

Sullivan, B.L., C.L. Wood, M.J. Iliff, R.E. Bonney, D. Fink, and S. Kelling. 2009. eBird: a citizen-based bird observation network in the biological sciences. Biological Conservation 142: 2282–2292.

Tacha, T.C., and Braun, C.E., editors. 1994. Migratory Shore and Upland Game Bird Management in North America. International Association of Fish and Wildlife Agencies, Washington, D.C.

Takats, D. L., Francis, C. M., Holroyd, G. L., Duncan, J. R., Mazur, K. M., Cannings, R. J., Harris, W., Holt, D. 2001. Guidelines for Nocturnal Owl Monitoring in North America. Beaverhill Bird Observatory and Bird Studies Canada, Edmonton, Alberta. 32 pp. http://www.birdscanada.org/download/owlguidelines.pdf.

Taylor, P. D., Crewe, T. L., Mackenzie, S. A., Lepage, D., Aubry, Y., Crysler, Z., Finney, G., Francis, C. M., Guglielmo, C. G., Hamilton, D. J., Holberton, R. L., Loring, P. H., Mitchell, G. W., Norris, D., Paquet, J., Ronconi, R. A., Smetzer, J., Smith, P. A., Welch, L. J., Woodworth, B. K.. 2017. The Motus Wildlife Tracking System: a collaborative research network to enhance the understanding of wildlife movement. - Avian. Cons. Ecol.12(1):8. https://doi.org/10.5751/ACE-00953-120108

Tozer, D. C. 2013. The Great Lakes Marsh Monitoring Program 1995-2012, 18 years of surveying birds and frogs as indicators of ecosystem health. Published by Bird Studies Canada, Port Rowan, ON. http://www.birdscanada.org/download/GLMMPreport.pdf.

Tozer, D. C. 2016. Marsh bird occupancy dynamics, trends, and conservation in the southern Great Lakes basin: 1996 to 2013. Journal of Great Lakes Research 42:136–145.

Tozer, D. C., Drake, K. L., Falconer, C. M. 2016. Modeling detection probability to improve marsh bird surveys in souther canada and the great lakes states. - Avian. Cons. Ecol. 11:3.

Taylor, C. M., and Norris, D. R. 2010. Population dynamics in migratory networks. - Theor. Eco. 3: 65–73.

US Fish and Wildlife Service. 2017. Waterfowl hunting management in North America Flyways.US: A collaborative effort of waterfowl managers across the continent. US Fish and Wildlife Service. http://flyways.us/ (Accessed 25 January 2017).

Van Wilgenburg, S. L., and Hobson, K. A. 2011. Combining stable-isotope (deltaD) and band recovery data to improve probabilistic assignment of migratory birds to origin. - Ecol. Appl. 21: 1340–51.

Wassenaar, L. I., and Hobson, K. A. 2003. Comparative equilibration and online technique for determination of non-exchangable hydrogen of keratins for use in animal migration studies. - Isot. Environ. Healt. S. 39: 211–217.

Webster, M. S., Marra, P. P., Haig, S. M., Bensch, S. and Holmes, R. T. 2002. Links between worlds: unraveling migratory connectivity. - Trends Ecol. Evol. 17: 76–83.

Yackulic, C.B., Chandler, R., Zipkin, E.F., Royle, J.A., James, D., Grant, E. H. C. & Veran, S. 2012. Presence-only modeling using MAXENT: when can we trust the inferences? - Meth. Eco. Evol. 4: 236–243

